# Rescue of normal excitability in LGI1-deficient epileptic neurons

**DOI:** 10.1101/2023.01.25.525523

**Authors:** Johanna Extrémet, Jorge Ramirez-Franco, Laure Fronzaroli-Molinieres, Norah Boumedine-Guignon, Norbert Ankri, Oussama El Far, Juan José Garrido, Dominique Debanne, Michaël Russier

## Abstract

Leucine-rich Glioma Inactivated 1 (LGI1) is a glycoprotein secreted by neurons, the deletion of which leads to Autosomal Dominant Lateral Temporal Lobe Epilepsy. Recently, we showed that LGI1 deficiency in a mouse model (KO-Lgi1) decreased Kv1.1 channel density at the axon initial segment (AIS) and at presynaptic terminals, thus enhancing both intrinsic excitability and glutamate release. However, the precise conditions for rescuing normal excitability in KO-Lgi1 neurons have still not been reported. Here we show that the selective expression of LGI1 in KO-Lgi1 neurons with the use of single-cell electroporation reduces intrinsic excitability, and restores both the Kv1.1 mediated D-type current and Kv1.1 immunostaining at the AIS. In addition, we show that the homeostatic shortening of the AIS length observed in KO-Lgi1 neurons is prevented in neurons electroporated with the Lgi1 gene. Furthermore, we reveal a spatial gradient of both intrinsic excitability and Kv1.1 immunostaining that is centred on the electroporated neuron. We conclude that expression of LGI1 restores normal excitability through the expression of functional Kv1 channels at the AIS.

## Introduction

LGI1 (Leucine-rich glioma inactivated) composed of a Leucine Rich Repeat (LRR) domain and an Epitempin (EPTP) domain is a secreted protein expressed primarily in the central nervous system (CNS). Several genes coding for proteins with EPTP domain are found in genomic region associated with epilepsy (Staub et al., 2002). The deletion of LGI1 protein induces Autosomal Dominant Lateral and Temporal Lobe Epilepsy (ADLTE), a pathology characterised by epileptic seizures with auditory disorders (Kalachikov et al., 2002). ADLTE can be induced by homozygous deletion of LGI1 in mice (Lgi1 Knock-Out mice or KO-Lgi1 mice). LGI1 functionality requires the binding to the disintegrin and metalloproteinase domain-containing protein 22 (ADAM22), a non-catalytic metalloprotease-like protein which act as a receptor for LGI1. Most of Lgi1 mutations prevent LGI1 secretion while secretion permissive mutations have been shown to disrupt its interaction with ADAM22 (Yokoi et al., 2015; Yamagata et al., 2018). Subsequent loss of LGI1 function in these mice induces epileptiform activities, seizures and premature death (Chabrol et al., 2010).

The mechanisms by which the epileptic phenotype occurs in KO-Lgi1 mice involve changes in excitatory synaptic transmission (Fukata et al., 2006, 2010; Lovero et al., 2015; Boillot et al., 2016), synapse maturation (Zhou et al., 2009; Lovero et al., 2015; Thomas et al., 2018), and also changes in intrinsic excitability (Schulte et al., 2006; Seagar et al., 2017; Lugarà et al., 2020). In Kv1.4/Kv1.1 channels, LGI1 prevents N-type inactivation by the Kvβ1 subunit (Schulte et al., 2006). Addition of recombinant LGI1 decreases intrinsic excitability in rat CA3 neurons whereas CA3 neurons of KO-Lgi1 mice are more excitable (Seagar et al., 2017). In fact, LGI1 tunes neuronal excitability by controlling the expression of voltage-gated Kv1.1-containing potassium channels density at the axon initial segment (AIS) of CA3 pyramidal neurons (Seagar et al., 2017). Accordingly, LGI1 and its receptor ADAM22 are colocalized at the AIS of dissociated hippocampal neurons, thus affecting the Kv1 potassium channel localization in this compartment (Hivert et al., 2019). The slowly inactivating D-type current (I_D_) mediated by Kv1 channels dampens action potential firing, and subsequently reduces synchronization of CA3 pyramidal neurons (Cudmore et al., 2010). Moreover, Kv1.1-containing channels present in the AIS have been shown to regulate intrinsic excitability of CA3 neurons (Rama et al., 2017). Consistent with a decrease in Kv1.1 density in CA3 neuron of KO-Lgi1 mice, the conductance of the voltage-gated D-type potassium current carried by the Kv1 channels family was found to be reduced (Seagar et al., 2017).

While rescue experiments have been often carried out to study synaptic mechanisms (Fukata et al., 2010; Lovero et al., 2015), the rescue strategy has not been used to confirm the role of LGI1 in the control of intrinsic excitability. In particular, it is unclear whether the rescue of Lgi1 gene in a single neuron is able to restore D-type potassium conductance and thus normal intrinsic excitability. We show here that selective expression of LGI1 in neurons from KO-Lgi1 mice using single-cell electroporation reduces intrinsic excitability and increases both the Kv1.1 mediated D-type current and Kv1.1 immunostaining at the AIS. In addition, we reveal a spatial gradient of both intrinsic excitability and Kv1.1 immunostaining that is centred on the electroporated neuron. Thus, the expression of LGI1 restores normal excitability through the expression of functional Kv1 channels at the AIS.

## Results

### Reduced excitability of KO-Lgi1 neurons electroporated with Lgi1-expressing plasmid

To investigate the involvement of LGI1 in shaping neuronal excitability, we rescued its expression in selected CA3 neurons from KO-Lgi1 mice organotypic slice cultures using single cell electroporation of an Lgi1-encoding plasmid (**Figure 1A**). A vector allowing expression of the enhanced green-fluorescence protein (EGFP) was used as control to confirm that changes in intrinsic excitability were not due to the single cell electroporation procedure itself (KO/EGFP neurons). CA3 pyramidal neurons were recorded in current-clamp and the number of action potentials for each increment of injected current was counted to establish input-output curves. Excitability parameters (first spike latency, rheobase and gain) were measured. No significant difference was observed between KO neurons and KO/EGFP neurons (**Figure 1B**) for the first spike latency (671 ± 54 ms n = 22 vs. 630 ± 50 ms n = 17, p=0.64; **Figure 1C**), for the rheobase (81 ± 9 pA n = 22 vs. 75 ± 11 pA n = 17, p = 0.65; **Figure 1C**), nor for the gain (0.11 ± 0.012 n = 22 vs. 0.12 ± 0.008 n = 17, p = 0.22; **Figure 1C**). We therefore chose KO/EGFP neurons as a control to describe our next results.

**Figure 1.**
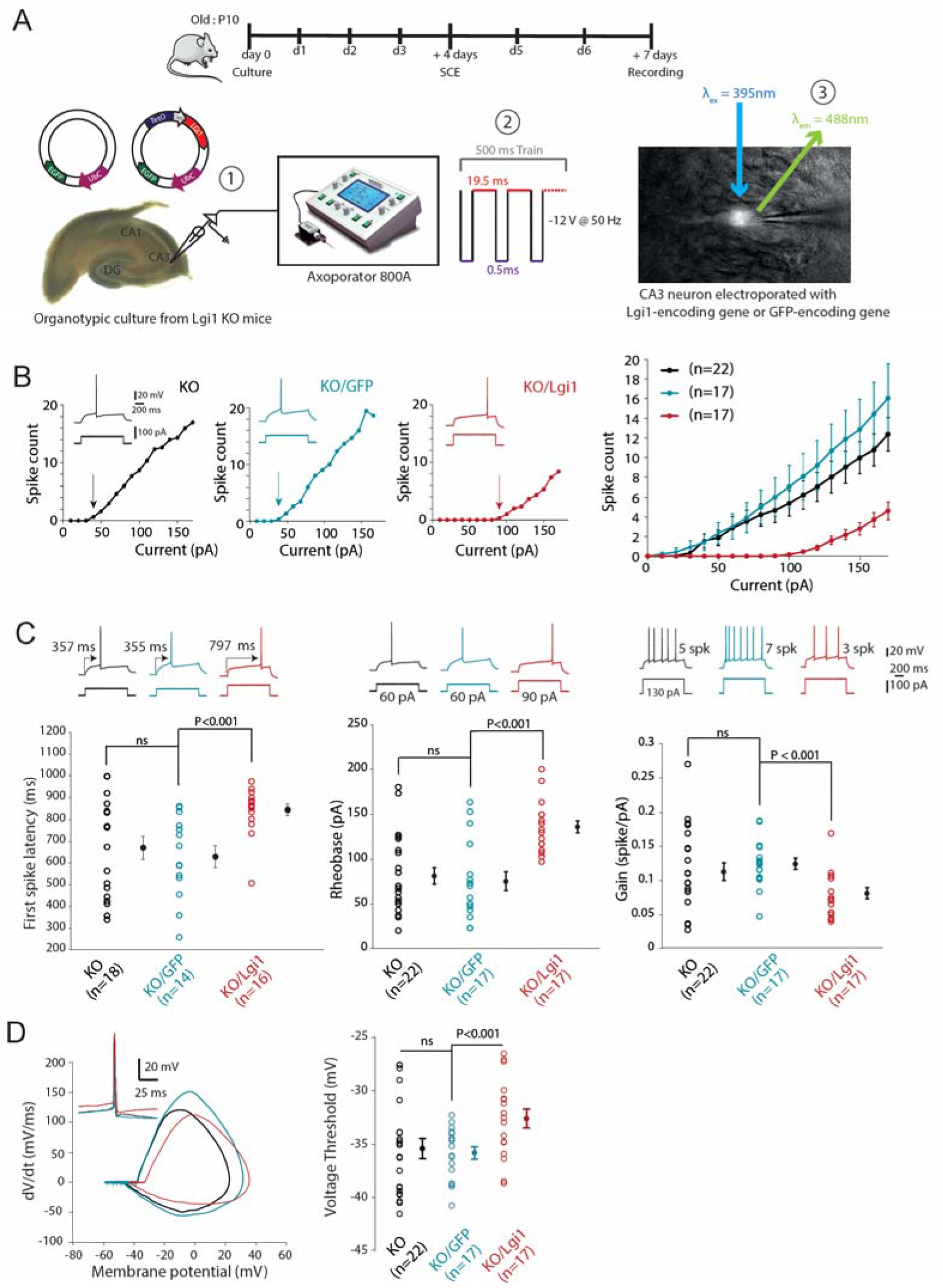
Rescue of intrinsic neuronal excitability in hippocampal CA3 pyramidal neurons from Lgi1 KO mice using SCE of an Lgi1-encoding plasmid. (A) Scheme of single-cell electroporation procedure. Top, the time course of the experiments (d0…d7 = age of organotypic slices in culture), (1) simplified representation of plasmid DNA constructs (UbC = eGFP constitutive promoter), (2) SCE protocol and (3) resulting fluorescent electroporated cell. λex = excitation wavelength, λem = emission wavelength. (B) Input-output curves showing the number of spike according to depolarising current increments and corresponding traces of representative neurons from organotypic culture of Lgi1 KO mice (black), electroporated with EGFP (blue) or Lgi1-encoding plasmids (red) respectively. Arrows indicate the rheobase. Right, averaged input-output curves. (C) Left, first spike latency obtained at rheobase current. Middle, rheobase. Right, gain. (D) Changes in spike threshold. Left, phase plot of action potentials recorded in a KO neuron (black), electroporated with EGFP (blue) and Lgi1 (red). Right, pooled data. Error bars represent SEM. Statistical analysis was performed using the Mann-Whitney test and significance was obtained at P<0.05.

In contrast to KO/EGFP neurons, KO neurons transfected with Lgi1 encoding plasmid (KO/Lgi1 neurons) expressed a clear ramp-and-delay phenotype (**Figure 1B**). This feature resembles the phenotype observed in WT CA3 neurons (Cudmore et al., 2010; Seagar et al., 2017). According to this characteristic, we measured an increase in the latency of the first spike in CA3 KO/Lgi1 neurons compared to KO/EGFP neurons (630 ± 50 ms, n = 17 in control vs. 844 ± 27 ms n = 17 in neurons electroporated with Lgi1-encoding plasmid, p < 0.001; **Figure 1C**).

Strikingly, the increased first spike latency in CA3 KO/Lgi1 neurons was paralleled with modifications in other excitability parameters. In fact, compared to control KO/EGFP neurons, the input-output curve of CA3 KO/Lgi1 neurons displayed a rightward shift (**Figure 1B**) with a significant increase in the rheobase (75 ± 11 pA n = 17 vs. 136 ± 7 pA n = 17, p < 0.001; **Figure 1C**), and a significant reduction in the gain (0.12 ± 0.008 n = 17 vs. 0.08 ± 0.008 n = 17, p < 0.001; **Figure 1C**). Furthermore, the voltage threshold of the action potential was found to be significantly depolarized in KO/Lgi1 neurons compared to KO or KO/EGFP neurons (−33 ± 0.5 mV, n = 17 in KO/Lgi1 neurons vs. -36 ± 0.5 mV, n = 17 in KO/EGFP neurons and -35 ± 0.5 mV, n = 22 in KO neurons; **Figure 1D**). Taken together, these data indicate that intrinsic excitability is reduced in CA3 KO/Lgi1 neurons compared to KO/EGFP.

### Increased sensitivity to DTx-k in KO/Lgi1 neurons

The ramp-and-delay phenotype observed before the evoked spike is a hallmark of slow inactivating D-type current (Cudmore et al., 2010). As this feature was observed in WT and KO/Lgi1 neurons, we examined the contribution of the D-type current to the electrophysiological phenotype after electroporation, by measuring the sensitivity to DTx-k in current-clamp. Only a slight difference was observed in the depolarization preceding the evoked spike (**Figure 2A**) with no significant difference in the first spike latency in KO/EGFP neurons after DTx-k application (n = 6, p = 0.52; **Figure 2C)**. In contrast, in KO/Lgi1 neurons, bath application of DTx-k induced a loss of the ramp-and-delay phenotype (**Figure 2B**) and a significant decrease of the latency to the first spike (n = 6, p = 0.03; **Figure 2C**). According to the input-output curves (**Figure 2A**), a slight increase in intrinsic excitability was noticed in KO/EGFP neurons after application of DTx-k. Clearly, the input-output curve of KO/Lgi1 neuron showed a more robust increase of intrinsic excitability after adding DTx-k (**Figure 2B**). While DTx-k did not induce changes in the rheobase of KO/EGFP neurons (n = 6, p = 0.44; **Figure 2C**), the rheobase of KO/Lgi1 neurons was significantly decreased after DTx-k application (n = 6, p = 0.03; **Figure 2C**). To quantify this effect, we subtracted the rheobase measured after application of DTx-k to the rheobase before adding DTx-k. Compared to the control, a fourfold increase of this rheobase difference was observed in KO/Lgi1 neurons (16.7 ± 6.8 pA, n = 6 vs. 67.2 ± 8.7 pA, n = 6, p = 0.004; **Suppl. Figure 1**) confirming a higher sensitivity to DTx-k in neuron rescued with LGI1.

**Figure 2.**
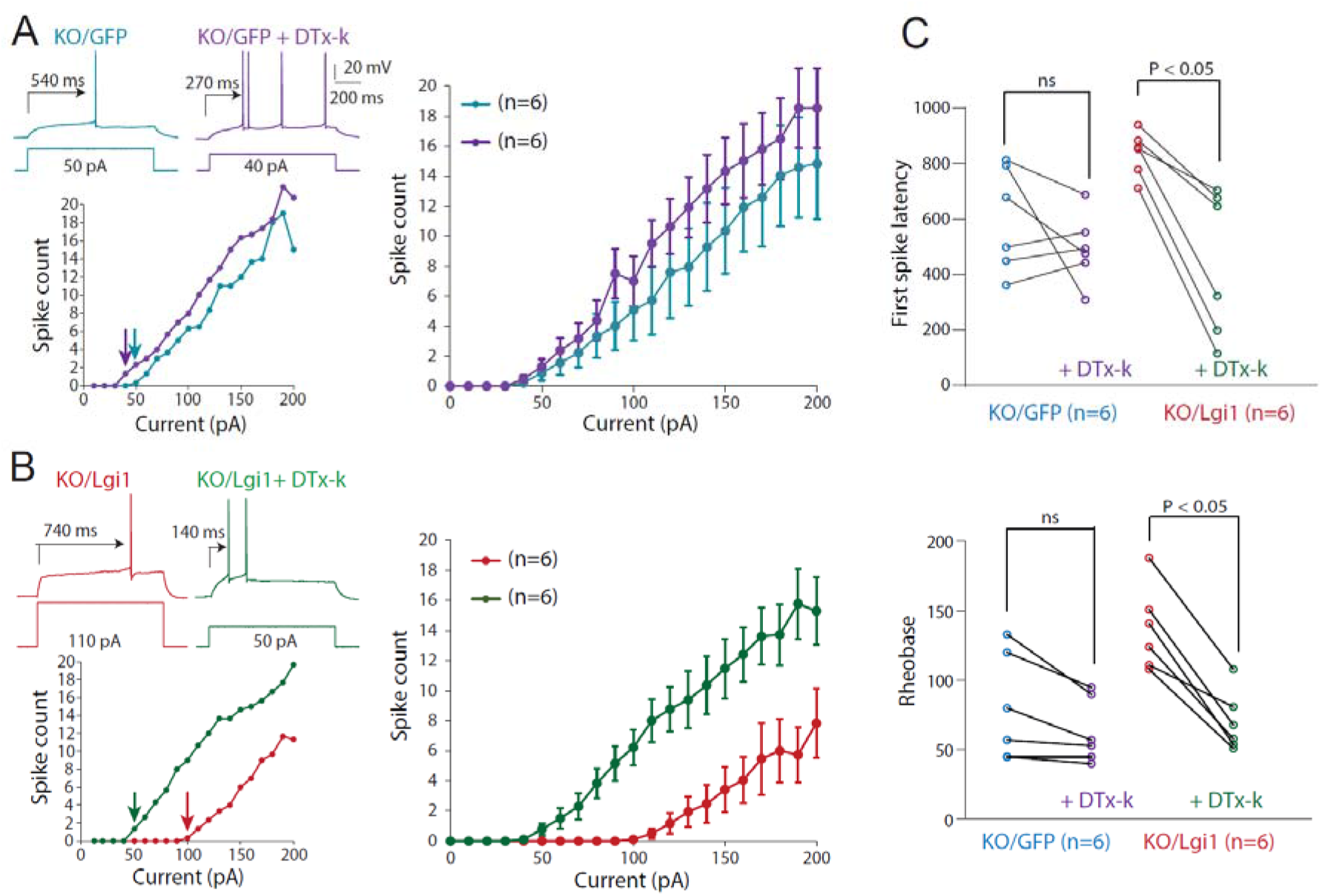
Rescued sensitivity to DTx-k in CA3 neurons electroporated with Lgi1 gene. (A) left, Input-output curves with corresponding traces of KO/EGFP (blue) representative neuron. DTx-k (100 nM) was bath applied on KO/EGFP neurons (purple). Rheobase is indicated on graphs by arrows. Right, Averaged input-output curves before and after bath application of DTx-k were obtained. (B) left, Input-output curves and corresponding traces of KO/Lgi1 representative neurons and right, averaged input-output curves between (red) and after (green) bath application of DTx-k. (C) Top, first spike latency obtained at rheobase current. Bottom, rheobase changes. Error bars represent SEM. Statistical analysis was performed using the Wilcoxon rank-signed test.

### Recovery of the D-type current and Kv1.1 channels at the AIS of KO/Lgi1 neurons

To confirm the recovery of D-type current in KO/Lgi1, we recorded neurons in voltage-clamp to obtain D-type conductance. For the same voltage step, the outward DTx-k-sensitive current evoked was weak in KO/EGFP neurons but highly increased in KO/Lgi1 neurons (**Figure 3A**). The D-type conductance was more than twofold in KO/Lgi1 neurons compared to the control (1.1 ± 0.4 nS, n = 6 in KO/EGFP neurons vs. 2.8 ± 0.5 nS, n = 8 in KO/Lgi1 neurons, p = 0.029; **Figure 3B**). The value of DTx-k-sensitive conductance reported in KO/lgi1 neurons was similar to that found in WT neurons (WT = 2.5 ± 0.2 nS, n = 15) (Seagar et al., 2017).

**Figure 3.**
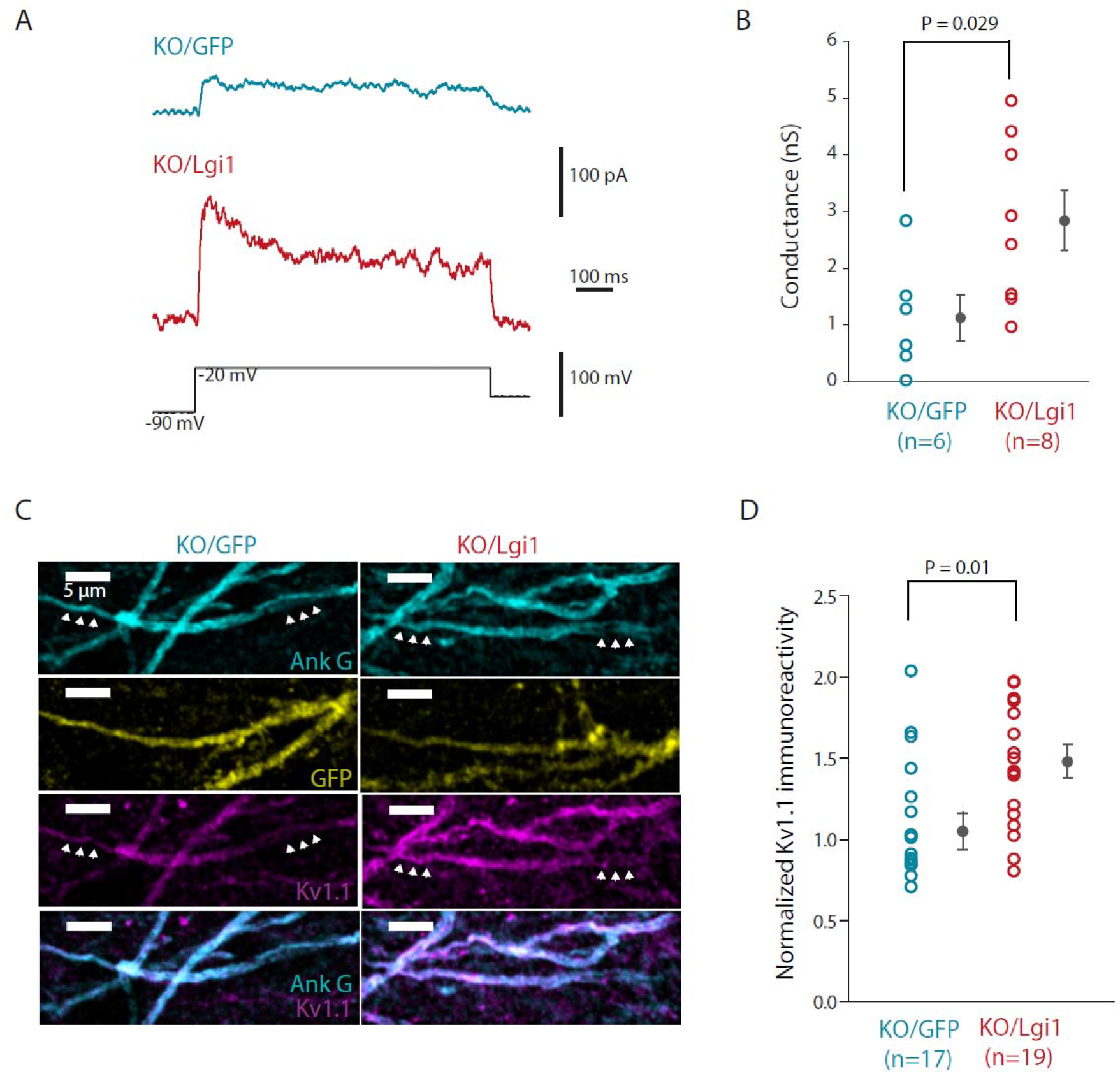
Rescue of the D-type current in CA3 neurons electroporated with Lgi1-expressing plasmid. (A) Representative traces of D-type current evoked by a voltage step from -90mV to -20 mV in CA3 KO/EGFP (top, blue) and KO/Lgi1 (bottom, red) neurons. (B) D-type conductance obtained in these two conditions. (C) Immunostaining of Ank G (cyan), EGFP (yellow) and Kv1.1 (magenta) at the AIS of KO/EGFP and KO/Lgi1 represented in column. Overlapping of Ank G and Kv1.1 labelling appear in white on the merge images (last row). Arrows indicate both ends of the AIS to follow its trajectory. (D) Quantification of Kv1.1 immunoreactivity at the AIS of KO/EGFP and KO/Lgi1 neurons. Error bars represent SEM. Statistical analysis was performed using the Mann-Whitney test.

In KO neurons, Kv1.1-containing channels were shown to be depleted mainly in the AIS (Seagar et al., 2017). We then evaluated Kv1.1-containing channel expression at the ankyrin G (Ank G)-positive AIS of KO/EGFP neurons and KO/Lgi1 neurons (**Figure 3C, 3D)**. The Kv1.1 staining was colocalised with that of Ank G and was predominant at the AIS of KO/Lgi1 neuron but not in KO/EGFP neuron (**Figure 3C**). Overall, compared to KO/EGFP neurons, the normalized immunostaining of Kv1.1-containing channel at the AIS was significantly increased in KO/Lgi1 neurons (1.0 ± 0.1, n = 17 in KO/EGFP neurons vs. 1.4 ± 0.1, n = 19 in KO/Lgi1 neurons, p = 0.01; **Figure 3D**), confirming that Lgi1 expression in CA3 neurons induced a D-type conductance recovery through a Kv1.1-containing channel rescue at the AIS.

### Rescue of AIS length in KO/Lgi1 neurons

AIS length is subject to homeostatic regulation following activity deprivation or activity enhancement (Kuba et al., 2010; Jamann et al., 2021). AIS length measured by β4-spectrin immunostaining was found to be significantly shorter in KO-Lgi1 mice than in WT mice (22.6 ± 0.1 µm, n = 98 vs. 35.2 ± 0.2 µm, n = 72; Mann-Whitney, p < 0.0001; **Figure 4A**). This ∼45% shortening of the AIS length suggests that the excitability increase observed in neurons lacking LGI1 (Seagar et al., 2017) is somehow compensated by an homeostatic process that makes the AIS shorter. We therefore checked whether restoring Lgi1 expression in neurons KO-Lgi1 neurons had an effect on AIS length. AIS length was found to be increased by ∼27% in KO/Lgi1 neurons compared to KO/EGFP neurons (28.3 ± 1.9 µm vs. 22.3 ± 1.6 µm, n = 10, Mann-Whitney test, p = 0.03; **Figure 4B**). This result supports the view that Lgi1 rescue is sufficient to recover the length of the AIS in KO-Lgi1 mice (**Figure 4C**).

**Figure 4.**
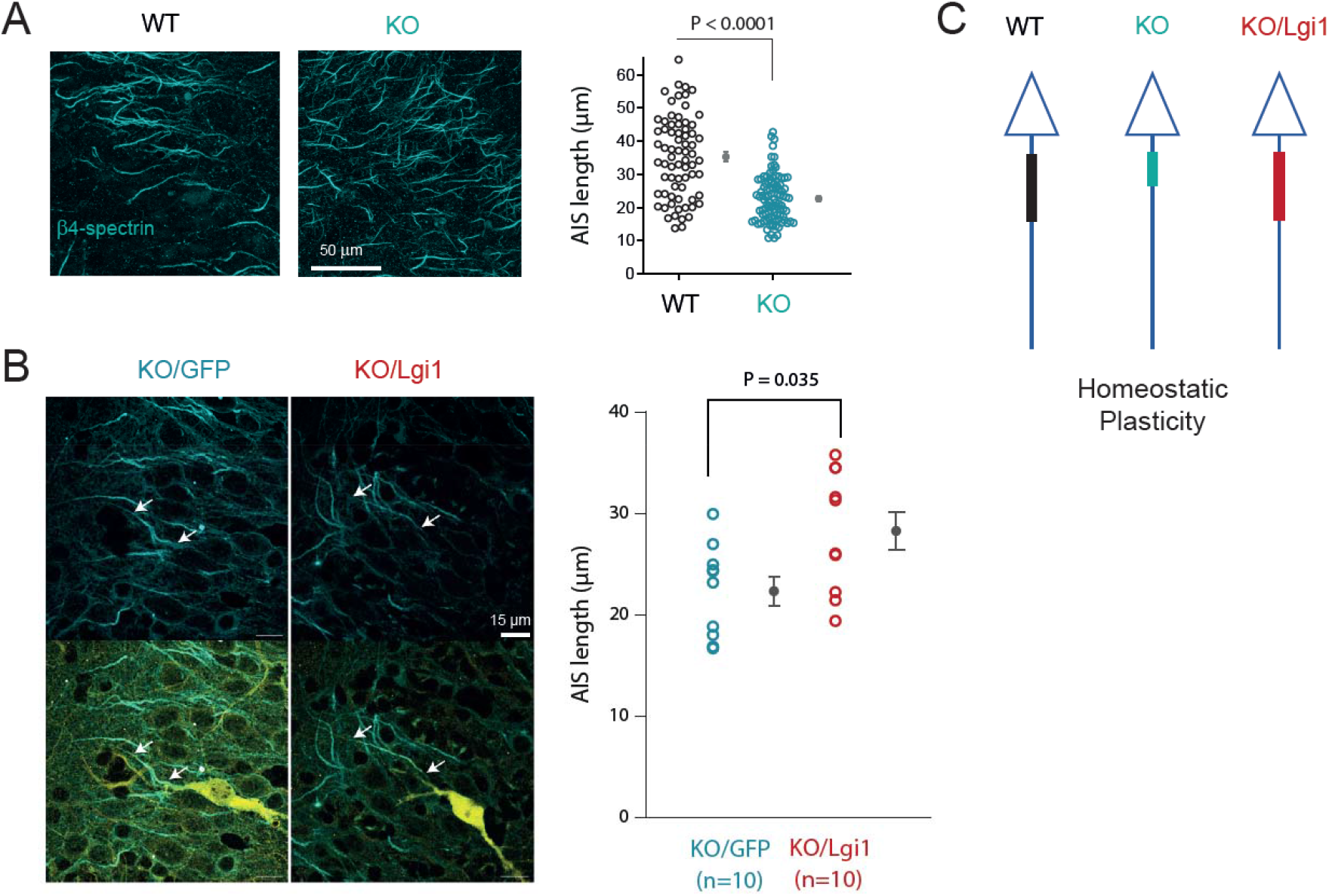
Rescue of AIS length in CA3 neurons electroporated with the Lgi1-expressing plasmid. A. Homeostatic reduction of the AIS length in Lgi1 KO mice. β4-spectrin immunolabeling in WT (left image) and in Lgi1 KO mice (right image). Right panel, group data (each dot represents one AIS). B. Comparison of AIS length in KO neuron electroporated with GFP (left column) and with Lgi1 (right column). Note the elongation of the AIS in a KO neuron electroporated with Lgi1. Right panel, group data. C. Summary scheme of the changes in AIS length as a function of Lgi1 expression.

### Modulation of the excitability of adjacent neurons by LGI1 spill-over

As LGI1 can be secreted by neurons (Senechal et al., 2005; Fukata et al., 2006; Lovero et al., 2015; Hivert et al., 2019), we checked the spatial extend of LGI1 rescue around an electroporated neuron. For this, CA3 neurons from KO-Lgi1 mice were electroporated with a Dendra2-tagged LGI1 (D2-LGI1) encoding plasmid (Ramirez-Franco et al., 2022). To reliably evaluate the extracellular labelling of D2-LGI1 protein, antibodies against Dendra2 were applied on living organotypic cultures before fixation and permeabilization. In these conditions, the D2-LGI1 protein immunostaining was extracellular, confirming the ability of LGI1 to be secreted by CA3 neurons (**Figure 5A**) (Ramirez-Franco et al., 2022). D2-LGI1 was concentrated around the soma and mostly at the ankyrin G-positive AIS of the KO/D2-Lgi1 neuron. Moreover, we could follow the labelling of secreted D2-LGI1 protein all along the axon. The proximal basal and apical dendrites were also enriched in secreted D2-LGI1 so that dots of secreted D2-LGI1 form a cloud around the soma. No D2-LGI1 was detected around distant neurons more than 200 µm from the electroporated neuron. Remarkably, D2-LGI1 was also highly detected along the AIS of neurons adjacent to the AIS and basal dendrites of the KO/D2-Lgi1 neuron (**Figure 5B**). To confirm that adjacent neurons to the electroporated neuron (i.e. nearby neurons) were targeted by the D2-LGI1, D2-LGI1 immunostaining was quantified at their AIS. D2-LGI1 immunostaining at the AIS of nearby neurons was about 15% that of KO/D2-Lgi1 neuron (196.9 au in KO/D2-Lgi1 neurons vs. 32.3 au in nearby AIS; **Figure 5C**). However, in comparison to AIS of a distant neuron, D2-LGI1 immunostaining was more than 7 fold higher at the AIS of a nearby neuron overlapping the AIS of the KO/D2-Lgi1 neuron (4.3 au in distant neuron vs. 32.3 au in nearby AIS; **Figure 5C**).

**Figure 5.**
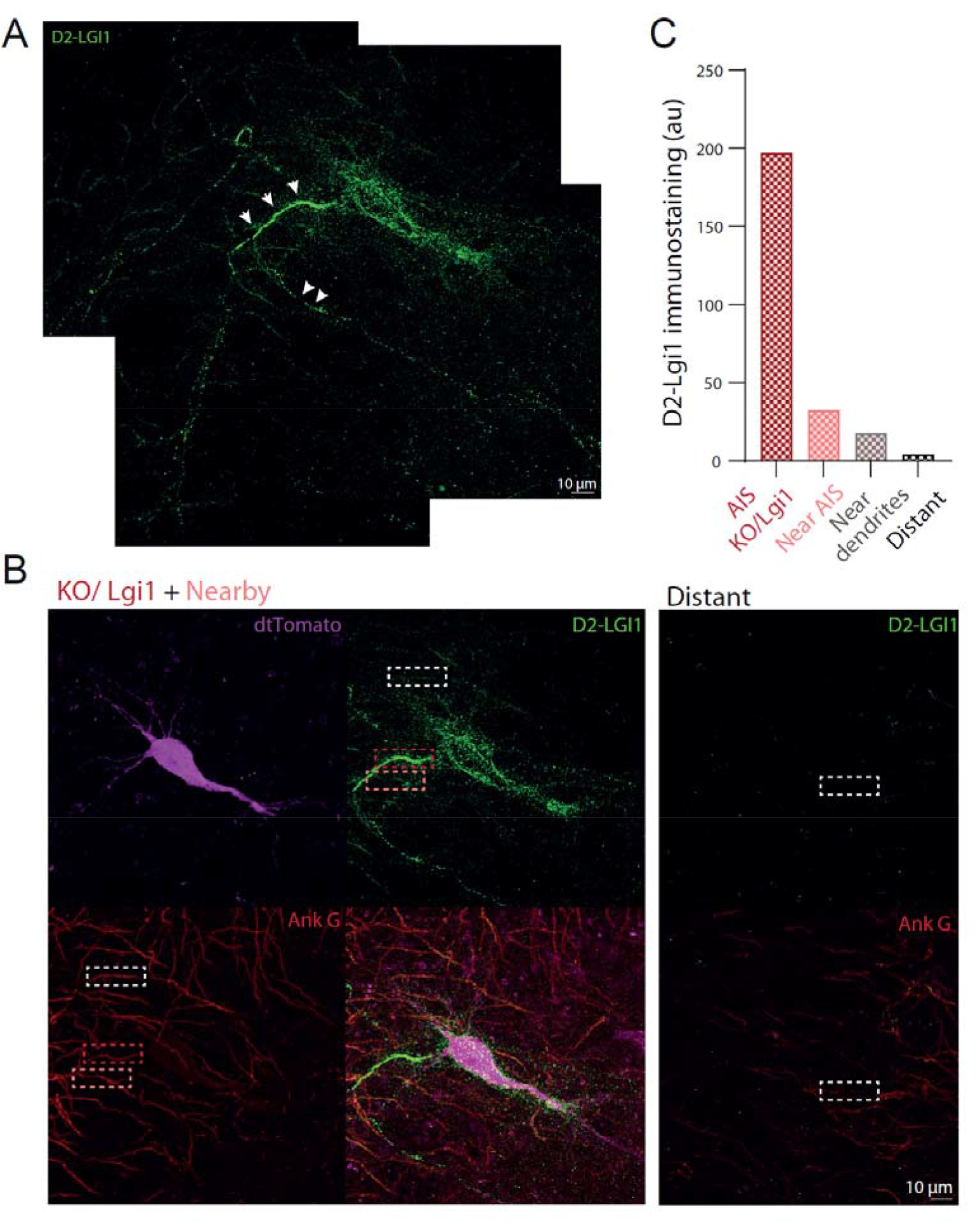
Spill-over of LGI1 on adjacent neurons. (A) Immunostaining of Dendra2(D2)-Lgi1 in KO neurons transfected with the D2-Lgi1. (B) dtTomato (magenta), Ank G (red) and D2-LGI1 (green) labelling are represented, with merge at the bottom corner for the field of view containing the KO/D2-Lgi1 neuron. A field of view at ∼200 µm away from the KO/D2-Lgi1 neuron (right). Dotted frames indicates AIS chosen for the quantification. (C) Quantification of the D2-LGI1 immunoreactivity at the AIS of the KO/D2-Lgi1 neuron (red), nearby KO/D2-Lgi1 AIS (pink), nearby KO/D2-Lgi1 dendrites (grey) and distant (black) neurons.

We then checked whether LGI1 extracellularly secreted by KO/Lgi1 neurons could affect the intrinsic excitability of nearby neurons (**Figure 6A**). Randomly chosen KO neurons were recorded in current clamp within a 150 µm wide field around the KO/Lgi1 neurons. KO neurons recorded at a distance of more than 200 µm from the KO/Lgi1 neurons were chosen as a control. Intrinsic excitability of these distant KO neurons was similar to that of KO neurons recorded in non-electroporated organotypic cultures (Rheobase: 81 ± 9 pA, n = 22 vs. 73 ± 3 pA, n = 22, p = 0.5; **Figure 1C** and **Figure 6A**). This confirms that neurons located far enough apart are not impacted by LGI1 secretion from the KO/Lgi1 neuron as suggested by the results of D2-LGI1 immunostaining. However, the input-output curve of nearby neurons presents a right shift that almost reach the curve of KO/Lgi1 neurons (**Figure 6A**). Interestingly, rheobase of nearby neurons was significantly increased compared to the distant KO neurons (100 ± 7.0 pA n = 25 vs. 73.4 ± 3.3 pA n = 22, p = 0.003; **Figure 6A**) but also significantly decreased compared to KO/Lgi1 neurons (100 ± 7.0 pA n = 25 vs. 129.0 ± 8.3 pA n = 29, p = 0.01; **Figure 6A**). This result indicates that secreted LGI1 is able to modulate intrinsic excitability of nearby neurons.

**Figure 6.**
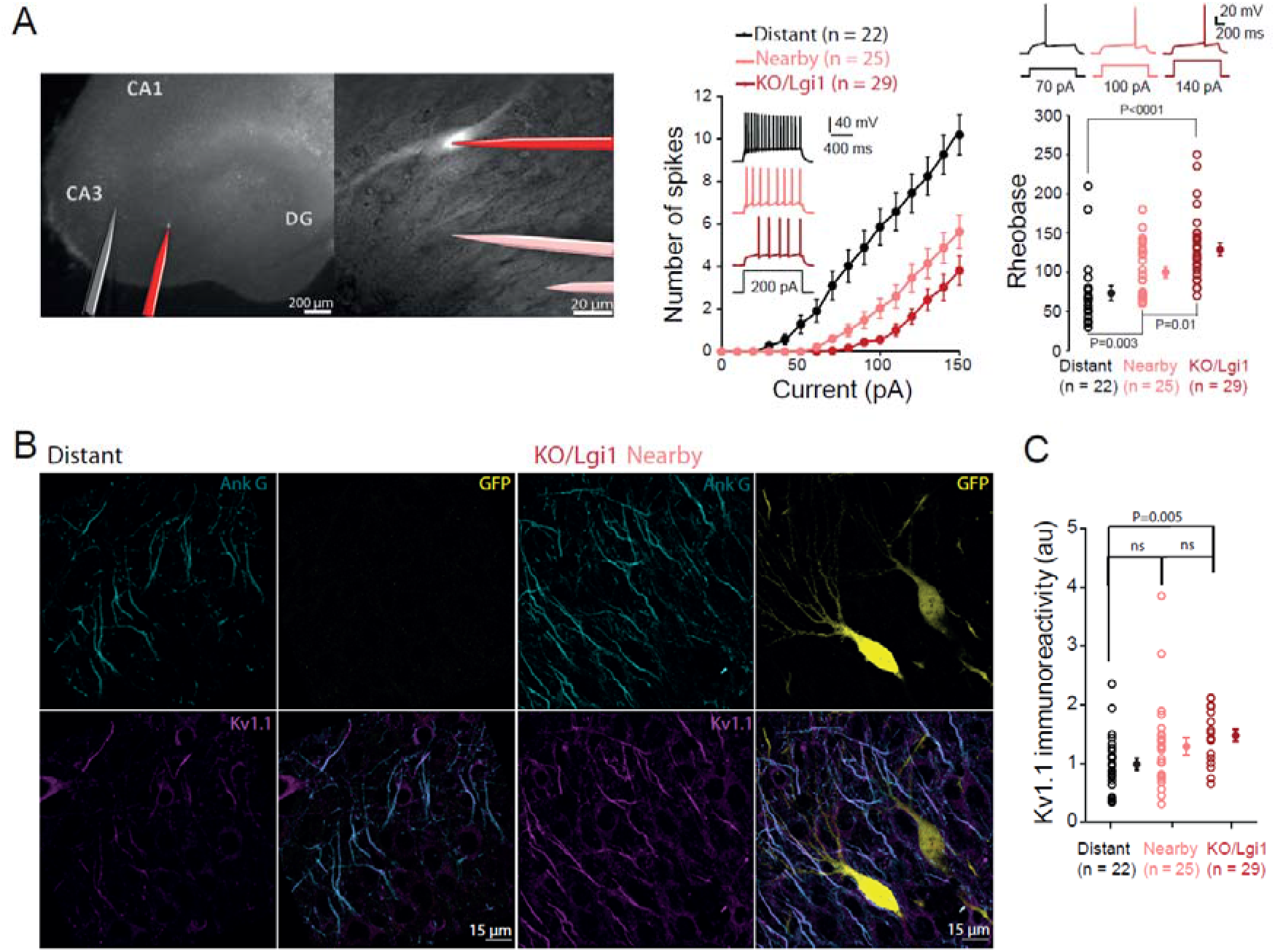
Paracrine reduction of intrinsic excitability induced by rescuing Lgi1 expression. (A) Hippocampal organotypic culture with a KO/Lgi1 neuron (left) and field of view of the KO/Lgi1 (right) with a scheme of the recording pipettes corresponding to distant (black), nearby (pink) and KO/Lgi1 (red) recorded neurons. Average input-output curves with corresponding traces and rheobase of the 3 conditions (bottom). (B) Immunostaining of Ank G (cyan), EGFP (yellow) and Kv1.1 (magenta) were represented in the field of view of distant and KO/Lgi1 neurons. (C) Quantification of Kv1.1 immunoreactivity at the AIS of distant, nearby and KO/Lgi1 neurons. Error bars represent SEM. Statistical analysis was performed using the Mann-Whitney in post-hoc test to Krukal-wallis test and significance was obtained at P<0.016.

As LGI1 protein rescued in KO/Lgi1 neurons allowed Kv1.1-containing channel recovery at the AIS, we measured Kv1.1 immunostaining at the AIS of nearby and distant neurons from KO/Lgi1 neurons (**Figure 6B**). As expected, Kv1.1 immunostaining was significantly increased in KO/Lgi1 neuron compared to controls (1.48 ± 0.10 au n = 19 vs. 0.99 ± 0.1 au; n = 25, p = 0.002; **Figure 6C**). However, despite a trends in Kv1.1 immunostaining from distant to nearby to KO/Lgi1 neurons, no significant difference was measured between nearby and distant neurons (1.29 ± 0.15 au n = 25 vs. 0.99 ± 0.10 au n = 25, p = 0.14; **Figure 6C**). In addition, no significant difference in Kv1.1 immunostaining was observed between nearby and KO/Lgi1 neurons (1.29 ± 0.15 au n = 25 vs. 1.48 ± 0.10 au n = 19, p = 0.07; **Figure 6C**). Our results thus indicate that the partial rescue of the rheobase in nearby neurons could be due to an elevated level of Kv1.1 channels at the AIS by ∼30%.

## Discussion

We show here that rescuing Lgi1 expression in KO-Lgi1 CA3 pyramidal neurons reduces their intrinsic neuronal excitability and increases both the DTx-k-sensitive D-type current and the density of Kv1.1 channels at the AIS. KO/Lgi1 neurons displayed a higher voltage AP threshold and a higher rheobase that was similar to that found in WT neurons (Seagar et al., 2017). In addition, the sensitivity to DTx-k that is absent in KO/EGFP neurons was rescued in KO/Lgi1 neurons. Furthermore, the AIS length shortening observed in KO neurons is partially suppressed in KO/Lgi1 neurons. As LGI1 is a secreted protein, we also addressed here the spatial extent of the changes in excitability and Kv1 channel expression using the labelling of LGI1 with D2. We reveal a gradient of both reduced excitability and of Kv1.1 channel recovery at the AIS of CA3 pyramidal neurons that is centred on the KO/Lgi1 neuron. Our results point to a major role of LGI1 in the control of Kv1.1 channels in the AIS of CA3 pyramidal neurons.

### LGI1 determines intrinsic excitability through Kv1.1 channels

So far, the effects of Lgi1 deletion have been extensively explored on both cellular excitability and Kv1 channel expression. In fact, a reduction in both the D-type current and Kv1.1 channels at the AIS has been reported in KO CA3 pyramidal neurons (Seagar et al., 2017). In KO-Lgi1 pyramidal neurons of the cortex, cellular excitability is increased due to a reduction in the number of Kv1.2 (Zhou et al., 2018). Furthermore, the reduction in Lgi1 expression using RNA silencing methods has also revealed an increase in neuronal excitability in dentate granule cells (Lugarà et al., 2020). We confirm here the link between LGI1 expression and Kv1 channel-dependent intrinsic excitability by showing that the rescue of Lgi1 expression reduces neuronal excitability through an elevation in the density of Kv1.1 channels and an enhancement of the DTx-k-sensitive D-type current.

Three lines of evidence support the fact that Kv1.1 channels are rescued in KO/Lgi1 neurons. First, the density of Kv1.1 channels was found to increase by 40% in KO/Lgi1 neurons. Second, the D-type current was increased by a factor 2 in KO/Lgi1. Third, the excitability was reduced (the spike threshold was elevated by ∼2 mV) and the sensitivity to DTx-k was restored. Taken together, our data demonstrate that LGI1 determines intrinsic excitability by controlling the expression of Kv1.1 channels at the AIS.

Kv1 channels are not only located at the AIS where they determine intrinsic excitability but they are also located at presynaptic terminals where they sharpen the spike and reduce transmitter release (Kole et al., 2007; Boudkkazi et al., 2011; Zbili et al., 2021). Interestingly, the up-regulation of intrinsic neuronal activity in KO-Lgi1 CA3 pyramidal cells comes with a loss of Kv1 channel function at the presynaptic terminal (Seagar et al., 2017). The effect of LGI1 rescue on synaptic function has not been tested in the present study.

### Rescue of AIS length

According to the homeostatic plasticity rules defined earlier (Turrigiano and Nelson, 2004; Grubb and Burrone, 2010; Kuba et al., 2010), the AIS length was found to be reduced by ∼45% in KO mice compared to the WT. Such shortening in AIS length has been already observed in status epilepticus induced with pilocarpine (Liu et al., 2017) but our study constitutes the first report of an AIS shortening in a genetic model of temporal lobe epilepsy. We show that rescuing the Lgi1 expression in LGI1-deficient neurons was sufficient to partially suppress the AIS length shortening (the mean AIS length was 28 µm in KO/Lgi1 neurons versus 35 µm in wild type neurons). Indeed, an increase by ∼27% was observed in electroporated neurons with the Lgi1-expressing plasmids. This partial rescue of the AIS length in KO/Lgi1 neurons is attributable to a network effect. Indeed, the CA3 area is supposed to remain epileptic or at least hyperexcitable despite the restoration of LGI1 protein in single CA3 neurons. This result also suggests that homeostatic plasticity of the AIS length depends on both intrinsic and local circuit factors.

### Spatial extent of paracrine release of LGI1

As LGI1 is a protein released by neurons, we used a D2-Lgi1 construct to show that LGI1 is mainly located at the AIS of the electroporated neuron and weakly present on the AIS of adjacent neurons. The excitability of neurons adjacent to the electroporated cell reveals a significant elevation of the rheobase in these neurons compared to that of more distant neurons. However, a non-significant increase in the Kv1.1 channel density (by ∼30%) was observed in KO neurons adjacent to KO/Lgi1 neurons, suggesting that electrophysiological analysis is more sensitive than immunostaining. Presynaptic and postsynaptic paracrine effects of LGI1 have been reported on synaptic transmission (Lovero et al., 2015). AMPA/NMDA ratio recorded in KO-Lgi1 neurons neighbouring a neuron transfected with Lgi1 was similar to WT neurons. Furthermore, the AMPA/NMDA ratio recorded in KO-Lgi1 CA1 neurons that receive inputs from CA3 neurons that were previously transfected with Lgi1-expressing lentivirus, was identical to that of WT neurons (Lovero et al., 2015). However, in this study the spatial extent of the paracrine release of LGI1 was not tested. In conclusion, the rescue of intrinsic excitability in KO-Lgi1 pyramidal neurons is mediated by an upregulation of Kv1 channels in the AIS and is tightly controlled by Lgi1.

## Methods

### Production of KO-Lgi1 Mice

All experiments were carried out according to the European and Institutional guidelines for the care and use of laboratory animals (Council Directive 86/609/EEC and French National Research Council) and approved by the local health authority (D13055-08, Préfecture des Bouches du Rhône). Heterozygous Lgi1^+/−^ mice were crossed to generate Lgi1^−/−^ (i.e. knock out, KO), Lgi1^+/−^, and Lgi1^+/+^ (wild type, WT) littermates. WT and KO mice were selected by analysing the amplification profile from Polymerase Chain Reaction (PCR). After sampling from 7 days old mice, DNA were denatured by temperature variations with a mix of 0.5 µl of DNA release diluted in 20 µl of dilution buffer (Master mix, ThermoFisher). 1 µl of DNA extraction from each sample was amplified with 0.25 µl of primer (final concentration of 0.3 µM for each primer: WT F, WT R, KO F, KO R; **Table 1**), 10 µl of Phire containing DNA polymerase (Master mix, ThermoFisher) and 8.75 of water in the Thermal Cycler (ProFlex PCR system, life technologies). PCR products were migrating in agarose gel (2%) and the resulting migration bands were visualised at 120 pb (WT band) and 200 pb (KO band) with a biomarker band as a reference.

**Table 1:**
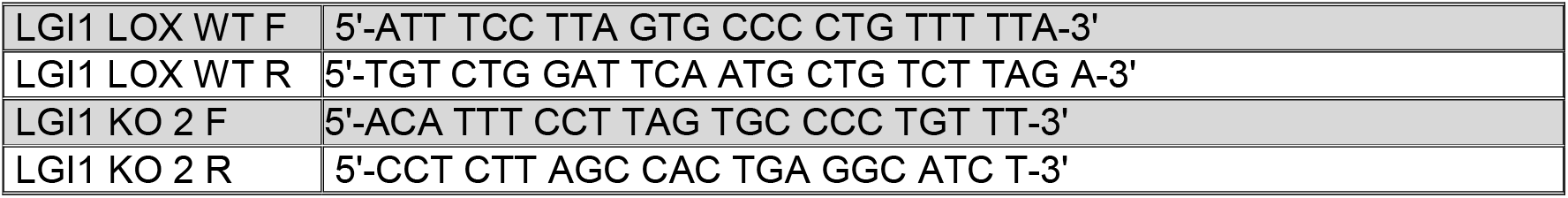

### Hippocampal slice cultures

Slices cultures were prepared as described previously (Debanne et al., 2008; Extrémet et al., 2022). Young Lgi1 KO mice (P7–P10) were killed by decapitation, the brain was removed, and each hippocampus dissected. Hippocampal slices (350 μm) were obtained using a Vibratome (Leica, VT1200S). They were placed on 20-mm latex membranes (Millicell) inserted into 35-mm Petri dishes containing 1 ml of culture medium and maintained for up to 21 d in an incubator at 34 °C, 95% O_2_–5% CO_2_. The culture medium contained (in ml) 25 MEM, 12.5 HBSS, 12.5 horse serum, 0.5 penicillin/streptomycin, 0.8 glucose (1 M), 0.1 ascorbic acid (1 mg/ml), 0.4 Hepes (1 M), 0.5 B27, and 8.95 sterile H_2_O.

### Single cell electroporation

Electroporation-mediated transfection (Rathenberg et al., 2003) was carried out in organotypic slice cultures of rat hippocampus at 4 days in vitro. The plasmid pIND-LGI1-IRES-EGFP (Ramirez-Franco et al., 2022) was used to concomitantly express Lgi1 and EGFP. In control conditions, we used pIND-IRES-EGFP (Ramirez-Franco et al., 2022) devoid of any Lgi1 expressing sequence. In order to follow extracellular LGI1 secretion, we used a pIND-Δ-IRES-Dendra2-LGI1 that expresses Dendra2-tagged Lgi1 (D2-Lgi1-encoding gene) (Ramirez-Franco et al., 2022). In order to identify the morphology of electroporated neurons, we used a tdTomato expressing plasmid (tdTomato-N1; gift from Michael Davidson & Nathan Shaner & Roger Tsien (Addgene plasmid # 54642 ; http://n2t.net/addgene:54642 ; RRID:Addgene_54642)) (Shaner et al., 2004). Before electroporation, plasmid solutions were centrifuged at 10000 g for 5 minutes to avoid obstruction of the micropipette. Immediately after single-cell electroporation, protein expression was induced by adding doxycycline (0.3 µg/ml) to the cell culture (**Figure 1A**). Lgi1 and EGFP were expressed for 3 days, and D2-Lgi1 and dtTomato were expressed for 6 days before further processing.

For single cell electroporation (SCE), the microscope chamber was consisting of a sterile 60-mm Petri dishes. The plasmid DNA constructs were diluted to a final concentration of 33 ng/µl in the internal solution containing (in mM): K-gluconate 120, KCl 20, Hepes 10, EGTA 0.5, MgCl_2_ 2, Na_2_ATP 2, and NaGTP 0.3 (pH 7.4). Micropipettes (7–10 MΩ) were filled with this DNA preparation after filtration with a sterile Acrodisc Syringe Filter (0.2 µm pore diameter).

During the SCE procedure, slice culture (DIV4-5) was positioned in the 60-mm Petri dishes and covered with pre-warmed and filtered external solution containing (in mM): 125 NaCl, 26 NaHCO_3_, 3 CaCl_2_, 2.5 KCl, 2 MgCl_2_, 0.8 NaH_2_PO_4_, and 10 D-glucose, equilibrated with 95% O_2_–5% CO_2_. The ground electrode and the microelectrode were connected to an isolated voltage stimulator (Axoporator 800A, Molecular Devices). Under visual guidance, the micropipette was positioned by a three-axis micromanipulator near the cell body of selected CA3 neurons. Pressure was controlled to have a loose-seal between the micropipette and the plasma membrane. When the resistance monitored reached 25 - 40 MΩ, we induced a train of -12V pulses during 500 ms (pulse width: 0.5 ms, frequency: 50 Hz). Each organotypic slice culture underwent SCE procedure for 8-10 selected neurons during a limited time of 30 min and was then back transferred to the incubator.

### Electrophysiology

After three days in the presence of doxycycline, whole-cell recordings were obtained from electroporated CA3 neurons, which were identified by GFP expression. Patch pipettes (7 – 10 MΩ) were filled with the internal solution containing (in mM): K-gluconate 120, KCl 20, Hepes 10, EGTA 0.5, MgCl_2_ 2, Na_2_ATP 2, and NaGTP 0.3 (pH 7.4). All recordings were made at 29 °C in a temperature-controlled recording chamber (Luigs & Neumann) perfused with the same external solution as for SCE. Liquid junction potential (∼-12 mV) was not compensated in the data reported. Neurons were recorded in current clamp or voltage clamp with a Multiclamp 700B Amplifier (Axon Instruments, Molecular Devices). Excitability was measured by delivering a range of long (1 s) depolarizing current pulses (10 – 250 pA, by increments of 10 pA) and counting the number of action potentials. Ionotropic glutamate and GABA receptors were blocked with 2 - 4 mM kynurenate and 100 μM picrotoxin respectively. Input-output curves were determined for each neuron and three parameters were examined: the rheobase (the minimal current eliciting at least one action potential), the gain (measured on each cell as the linear fit of the spike number as a function of current pulse) and the latency of the first spike (depolarising time before the evoked spike under rheobase current eliciting only 1 spike). Sensitivity to Kv1-channel blocker dendrotoxin K (DTx-K) was determined by current-clamp recording before and after bath application of the DTx-K (100 nM). Voltage-clamp protocols to measure Kv1-channel mediated currents consisted of a family of voltage-step commands from a holding potential of -90 mV (step from -80 to +10 mV in 10-mV increments). To block voltage dependent Ca^2+^ and Na^+^ currents, 200 μM Ni^2+^, 50 μM Cd^2+^, and 0.5 μM TTX were added to the extracellular solution. Algebraic isolation of the D-type potassium current was done by subtracting the currents evoked in the presence of the selective blocker of Kv1.1-containing channel DTx-K, from currents evoked in control solution. The D-type conductance was calculated from the maximal peak of the DTx-k-sensitive current obtained and the driving force of potassium in our recording conditions (electrochemical potential of potassium E_K+_=-105.09 mV). In all voltage-clamp experiments, leak and capacitance subtraction was performed using a p/n (n = 4) protocol. The voltage and current signals were low-pass filtered (3 - 0.4 kHz respectively), and acquisition was performed at 10 kHz with pClamp10 (Axon Instruments). Data were analysed with ClampFit (Axon Instruments) and Igor (Wavemetrics). Spike thresholds were measured using phase plots (Fékété et al., 2021).

### Immunohistochemistry

Organotypic cultures from KO-Lgi1 mice were performed as described previoulsy (Ramirez-Franco et al., 2022). Briefly, slices were fixed in a solution containing 4% of paraformaldehyde in PBS for 15 min at 4°C, incubated in 50 mM NH4Cl in PBS for 15 min at room temperature (RT), and blocked overnight at 4°C in a solution containing 5% Normal Goat Serum (NGS, Vector laboratories) and 0.5% Triton X-100 in PBS. After blocking, slices were incubated (24h; 4°C) with primary antibodies in a solution containing 0.5% triton X-100 and 2% NGS in PBS. The following antibodies were used: guinea pig anti-AnkG (1:400, Synaptic Systems, 386005, RRID: AB_2737033), rabbit anti-GFP (1:500, Synaptic Systems, 132003, RRID: AB_1834147), mouse anti-Kv1.1 (1:200, Antibodies Incorporated, 75-105, RRID: AB_10673166) or mouse anti-AnkG (1:400, Antibodies incorporated, 75-147, RRID: AB_10675130), rabbit anti-Dendra2 (1:200, Antibodies online, ABIN361314, RRID: AB_10789591), chicken anti-tdTomato (1:500, ThermoFisher, TA150089). The slices were then washed 4 times for 20 min each time in PBS 0.5% Triton X-100 and then incubated with the appropriate secondary antibodies for 2h at RT in a solution containing 0.5% Triton X-100 and 2%NGS in PBS. The following antibodies were used: Alexa Fluor 647 goat anti-guinea pig (1:150, Jackson Immunoresearch), Alexa Fluor 488 goat anti-rabbit (1:200, Jackson Immunoresearch), Alexa Fluor 594 goat anti-mouse (1:200, Jackson Immunoresearch) or Alexa Fluor 488 goat anti-rabbit (1:200, Jackson Immunoresearch), Alexa Fluor 594 donkey anti-chicken (1:400, Jackson Immunoresearch), Alexa Fluor 647 donkey anti-mouse (1:400, Molecular Probes). Subsequently, sections were washed 3 times for 20 min in PBS 0.5% Triton X-100. Nuclei were stained using DAPI at a final concentration of 1.5 µg/ml in PBS for 10 min and washed in PBS for 15 min. Slices were then mounted in Vectashield mounting medium (Vector laboratories). Z-stacks of confocal sections were acquired on a LSM780 confocal scanning microscope (Zeiss).

### Image processing and quantification

ImageJ (NIH) was used for image analysis and quantification. In order to measure Kv1.1 immunostaining at the AIS, rolling ball background subtraction was performed over the images. Then, object identification was done by using 3D object counter plugin in ImageJ over AnkG signal (Bolte and Cordelières, 2006). After object detection, this 3D mask was applied over Kv1.1 signal or D2-Lgi1 signal, and the 3D ROI manager (Ollion et al., 2013) was used to identify individual AISs within the field of view. Kv1.1 or D2-Lgi1 Immunostaining levels were collected as average grey values. For each condition, Kv1.1 immunostaining at the AIS was normalized to KO neurons away from the neuron electroporated with Lgi1.

In order to overcome differences in EGFP expression between different cells, and since EGFP was only used as a reporter to identify transfected cells, different minimum and maximum display settings were applied for the EGFP channel in some of the images.

AIS length was measured using β4-spectrin staining. Briefly, a line was drawn starting from each soma down to its axon following the β4-spectrin staining. This defines a fluorescence intensity profile along the AIS that increases and then decreases. Starting and end positions of the AIS are defined as the positions where β4-spectrin intensity is detectable.

### Statistics

Pooled data are presented as mean ± SEM and statistical analysis was performed using the Mann–Whitney U test or Wilcoxon rank signed test and Kruskal-Wallis test followed by Mann-Whitney post-hoc test for multi-comparison. Data were considered statistically significant for p ≤ 0.05 except for Mann-Whitney post-hoc test in which significant is reached at p ≤ 0.016 (0.05/3). For some conditions in electrophysiology and in immunohistochemistry, a resampling was carried out by using the bootstrap method to have a comparable workforce between distant KO neuron, nearby KO neuron, KO/Lgi1 neurons and KO/EGFP neurons.

## Author contributions

DD, OEF & MR designed research; JE, JRF, NBG, JJG & MR performed research; JE, JRF, NA, JJG, DD & MR analysed data; JE, DD & MR wrote the manuscript.

## Acknowledgments

Supported by INSERM, CNRS, Aix-Marseille Université, Agence Nationale de la Recherche (ANR-17-CE16-022-01 to DD and ANR-17-CE16-022-02 to OEF) and Fondation pour la Recherche Médicale (FRM DEQ20180839583 to DD).

---

**Supplementary Figure 1.**
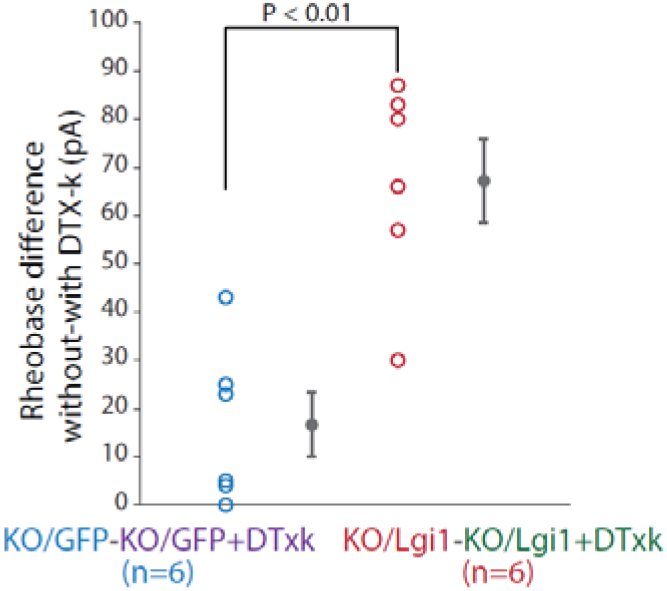
Effect of DTx-k on rheobase. The DTx-k effect was obtained by subtraction of the rheobase after adding DTx-k to the rheobase before DTx-k and compared between KO/EGFP and KO/Lgi1. Error bars represent SEM. Statistical analysis was performed using the Mann-Whitney test and significance was obtained at P<0.05.

